# Injury size regulates glucose allocation locally and systemically during vertebrate tissue regeneration

**DOI:** 10.64898/2026.08.31.748065

**Authors:** Claudia Kuntner, Cécile Philippe, Chrysoula Vraka, Lena Zachhuber, Thomas Wanek, Joachim Friske, Victoria Weissenböck, Thomas Helbich, Marcus Hacker, Elly M Tanaka, Leo Otsuki

## Abstract

Tissue regeneration requires careful allocation of metabolic resources, yet how organisms adjust this allocation in response to varying amounts of tissue loss remains poorly understood. Here, we show that the regenerative metabolic response is not fixed: the size of an injury regulates how glucose is allocated at both local and organism-wide levels. We first demonstrate that tail regeneration requires glucose metabolism in the axolotl (*Ambystoma mexicanum*), a salamander capable of regenerating centimetre-scale tissues. We then mapped glucose uptake in axolotls regenerating from small or large tail injuries using positron emission tomography/magnetic resonance imaging (PET/MRI) and the radiolabelled glucose analogue [^18^F]FDG. Glucose uptake was elevated in regenerating tails compared to uninjured tails. During early regeneration, larger injuries induced higher glucose uptake than smaller injuries, correlating with faster regenerative outgrowth. Larger injuries also increased glucose uptake in distant organs, indicating a systemic metabolic response. Together, our findings suggest that metabolic responses tuned to injury size underlie faithful tissue regeneration and establish PET/MRI as a powerful approach for studying whole-body metabolic dynamics in large regenerating vertebrates.

## Introduction

Regenerative animal models have revealed that tissue regeneration requires rewiring of cellular metabolism. Injury elevates glucose metabolism, which is required for regeneration of tail, fin and heart in zebrafish (*Danio rerio*) ^1–4^ and of the spinal cord and tail in *Xenopus* species tadpoles ^5,6^. This requirement is not explained solely by increased energy demand, as blocking oxidative phosphorylation does not inhibit regeneration ^2^. Glucose is preferentially shuttled into glycolysis, the pentose phosphate pathway or hexosamine biosynthesis, depending on the injury model. Although zebrafish and tadpoles are well-suited to microscopic imaging and high-throughput perturbations, their small size limits the study of larger injuries. This raises the question of whether similar principles apply to the regeneration of larger, centimetre-scale tissues.

Axolotls are aquatic vertebrates well-suited to studying centimetre-scale tissue regeneration. They regenerate throughout life while growing continuously in size, eventually exceeding 30 cm in length. Axolotls and other salamanders regenerate many body parts, including the tail, limbs, heart and parts of the nervous system ^7^, yet studies of metabolism have been scarce. Histochemical assays and metabolic profiling suggest a switch from oxidative phosphorylation towards aerobic glycolysis during salamander limb regeneration ^8–12^. Autoradiography demonstrated increased glucose and acetate uptake after cryoinjury to the axolotl heart ^13^. However, it remains unknown if glucose metabolism is functionally required for tissue regeneration in salamanders, and whether metabolic responses vary with the amount of tissue loss. Here, we address both questions using the axolotl tail model.

The axolotl tail contains tissues also present in humans, including the spinal cord, vertebrae and trunk muscle. It is well-suited to quantitative analyses due to its linear, segmentally repeating structure, which facilitates the removal of precise amounts of tissue and measurement of subsequent outgrowth ^14–16^. Tail amputation induces the formation of a proliferative cell mass, called a blastema, at the tail tip, which differentiates to restore the missing tissues. We tested whether glucose metabolism is required for the blastema to regenerate the axolotl tail, as in zebrafish and *Xenopus* species.

To identify tissues with higher glucose demand, we established protocols for simultaneous positron-emission tomography / magnetic resonance imaging (PET/MRI) in large (14 cm) regenerating axolotls. PET is a non-invasive biomedical imaging modality that enables the visualization of metabolic processes *in vivo* by detecting the distribution of radiolabelled tracers throughout the body. When combined with MRI, PET signals can be assigned to specific anatomical structures, enabling whole-body metabolic mapping in living organisms ^17^. One of the most widely used radiotracers is the radiolabelled glucose analogue 2-deoxy-2-[^18^F]fluoroglucose ([^18^F]FDG, hereafter “FDG”), used to evaluate glucose hypermetabolism (e.g. in cancer) or hypometabolism (e.g. in neurodegeneration). FDG is taken up into cells via GLUT 1 and GLUT 3 transporters but, unlike glucose, becomes trapped as FDG-6-phosphate after the first step of glycolysis. Hence, FDG-PET allows mapping of glucose uptake across the body, while MRI provides three-dimensional anatomical information. As PET/MRI does not require tissue fixation or light penetration, it enables the study of large, living animals that were previously inaccessible with conventional microscopy. MRI has previously been used to study the regenerating axolotl heart and brain ^18–20^. By contrast, PET has only been used in uninjured axolotls, for example to study the adrenal stress response ^21,22^, and in one review article presenting a pilot experiment on regenerating heart ^23^. Thus, we reasoned that there was great potential to harness PET for in-depth studies of vertebrate regeneration.

We first established a practical FDG-PET/MRI imaging workflow for axolotls, to advance the limited methodological details in this species during regeneration ^23^. We then mapped whole-body FDG uptake during early and mid-regeneration following either a large (4 cm) or small (2 cm) tail amputation. Our results demonstrate that both local and systemic changes in metabolism underlie injury size-matched tissue regeneration.

## Results

### Glucose metabolism is required for tail regeneration in axolotls

To test whether glucose metabolism is required for tail regeneration, we treated regenerating axolotls with 2-deoxy-D-glucose (2-DG). 2-DG is a non-metabolizable glucose analogue that is taken up by glucose-consuming cells *via* GLUT transporters and inhibits glycolysis and related pathways. We induced regeneration by removing 0.5 cm of the tail tip of 4 cm axolotls, then injected regenerating animals every 2 days with either 2-DG (test) or glucose (control) (**Figure 1A**). We measured the outgrowing spinal cord as a proxy for overall tail regeneration. Spinal cord grows out unidirectionally and can be labelled genetically using a *Sox2:mCherry* transgene ^24^.

**Figure 1.**
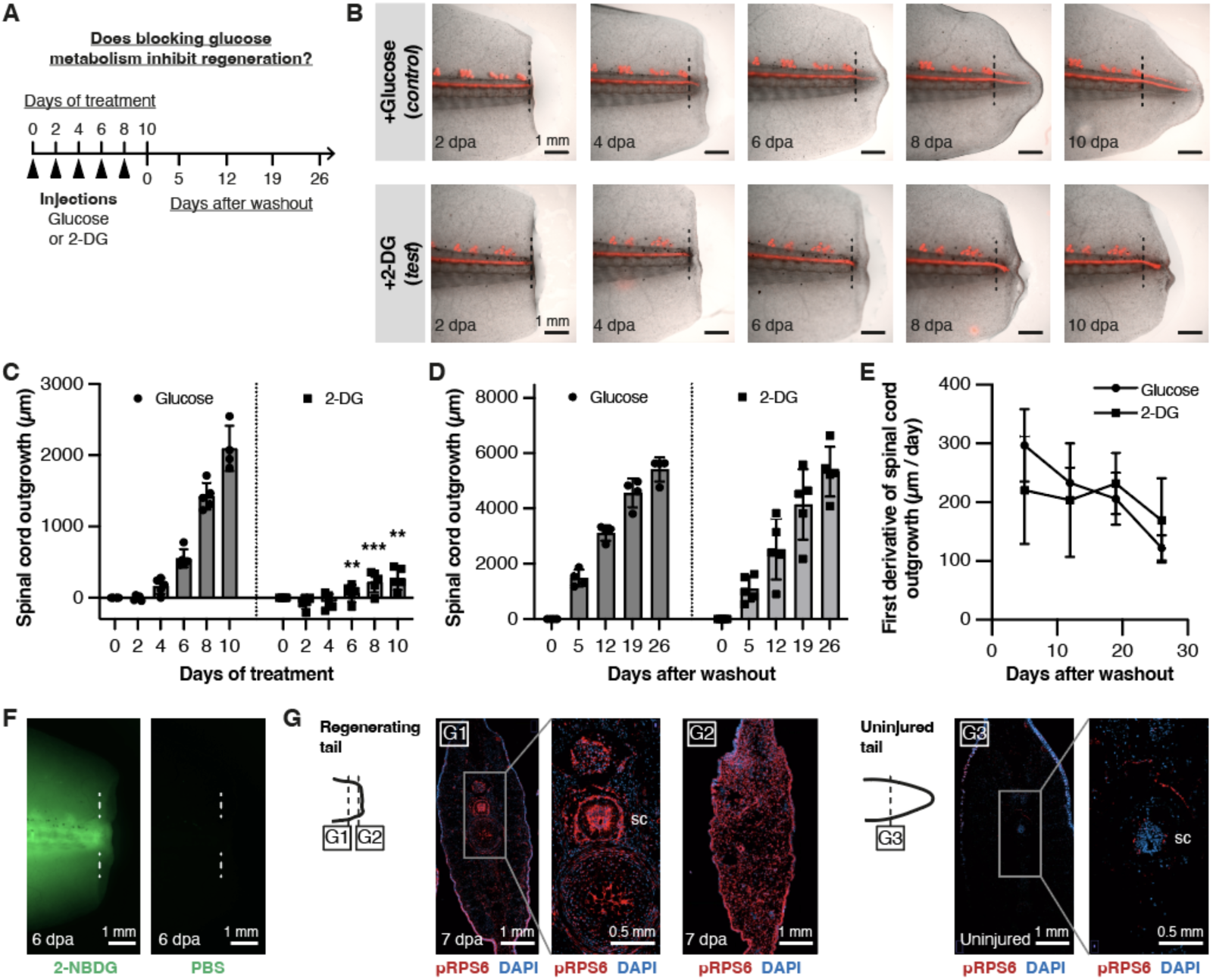
Tail regeneration requires glucose metabolism in axolotls. (A) Schematic of 2-DG experiments. Glucose (control) or 2-DG (test) was injected into axolotls every 2 days from day 0 until day 8 post-amputation (dpa). No further injections were performed from day 10 (= washout period). (B) Time series of tail regeneration in axolotls injected with glucose (top) or 2-DG (bottom). Dotted lines indicate amputation plane. *Sox2*:mCherry (red) labels cells in the spinal cord and lateral line. (C) Spinal cord outgrowth in axolotls treated with glucose (left) or 2-DG (right). 2-DG significantly reduces tail outgrowth compared to controls. From left to right, **: *p* = 1.50×10^-3^, ***: *p* = 2.88×10^-5^, **: *p* = 2.27×10^-3^, Šidák’s multiple comparisons. *n* = 5 animals per condition. (D) Spinal cord outgrowth after washout of glucose (left) or 2-DG (right). There is no significant difference between these conditions. 2-way ANOVA. *n* = 4 animals (control) or 5 animals (test). (E) Rate of spinal cord outgrowth after washout of 2-DG or glucose. Calculated from the data in (D). There is no significant difference between these conditions. 2-way ANOVA. (F) Widefield images of 6 dpa axolotls 2 hours after injection with 2-NBDG (fluorescent glucose analogue) or PBS (negative control). Dotted lines indicate amputation plane. (G) Single section confocal images of regenerating tail (G1-G2) or uninjured tail (G3), stained for pRPS6 (red) and DAPI (blue). G1 depicts stump tissue neighbouring the blastema, with the spinal cord indicated (sc). G2 depicts tail blastema. G3 depicts tissue from the equivalent region of an uninjured tail. Representative of *n* = 4 tails per condition.

2-DG significantly reduced spinal cord and overall tail outgrowth compared with controls as shown on day 6, 8 and 10 of treatment (*n* = 5 animals per group; **Figures 1B-C**). Thus, glucose metabolism is required for axolotl tail regeneration. An important question is whether cells are responsive to glucose only immediately after injury, or whether this sensitivity is sustained over long periods. When we suspended 2-DG treatment at 10 days post-amputation (10 dpa), the blocked spinal cords resumed outgrowth - and did so at rates indistinguishable from controls (**Figures 1D-E**). These results indicate that regenerative cells remain responsive to glucose for at least 10 days after injury.

### Injury induces metabolic activation in the regenerating tail tip

To locate the glucose-consuming cells in the tail, we pulsed regenerating axolotls with the fluorescent glucose analogue 2-(N-(7-Nitrobenz-2-oxa-1,3-diazol-4-yl)Amino)-2-Deoxyglucose (2-NBDG). Like 2-DG, 2-NBDG is taken up by cells and serves as a fluorescent readout for glucose uptake. We readily detected 2-NBDG signal in pulsed axolotls, but did not observe accumulation in the blastema, possibly due to high uptake by the neighbouring muscle and vertebrae (**Figure 1F**).

As an alternative assay, we collected tissue sections from uninjured or regenerating tails and performed immunostaining with an antibody targeting phosphorylated ribosomal protein S6 (pRPS6^Ser240/244^) ^25^. pRPS6 indicates activation of the mTORC1 pathway, which stimulates metabolic pathways including glycolysis and would be consistent with increased glucose consumption. pRPS6 was upregulated in blastemas and in neighbouring stump tissue compared to the equivalent region of uninjured tails (**Figure 1G**). pRPS6 was present throughout the tail blastema, except in putative blood cells, and throughout neighbouring stump tissue, particularly within the spinal cord. Within the spinal cord, both ependymoglia and neurons upregulated pRPS6 after injury, contrasting with the sparse signal in peripheral neurons in uninjured controls.

Together with the 2-DG experiments, our results support a model in which axolotls increase glucose uptake locally at the injury site, a requirement for regeneration.

### Extending metabolic analyses to large axolotls

Our 2-DG experiments were performed in small (4 cm) axolotls. We next asked whether similar metabolic activation occurs in larger (14 cm) axolotls regenerating several centimetres of tissue (**Figure 2A**). In contrast to small animals, larger axolotls are difficult to image live using conventional 3D optical approaches as their increased tissue thickness limits light penetration, necessitating tissue clearing (**Figure 2B**).

**Figure 2.**
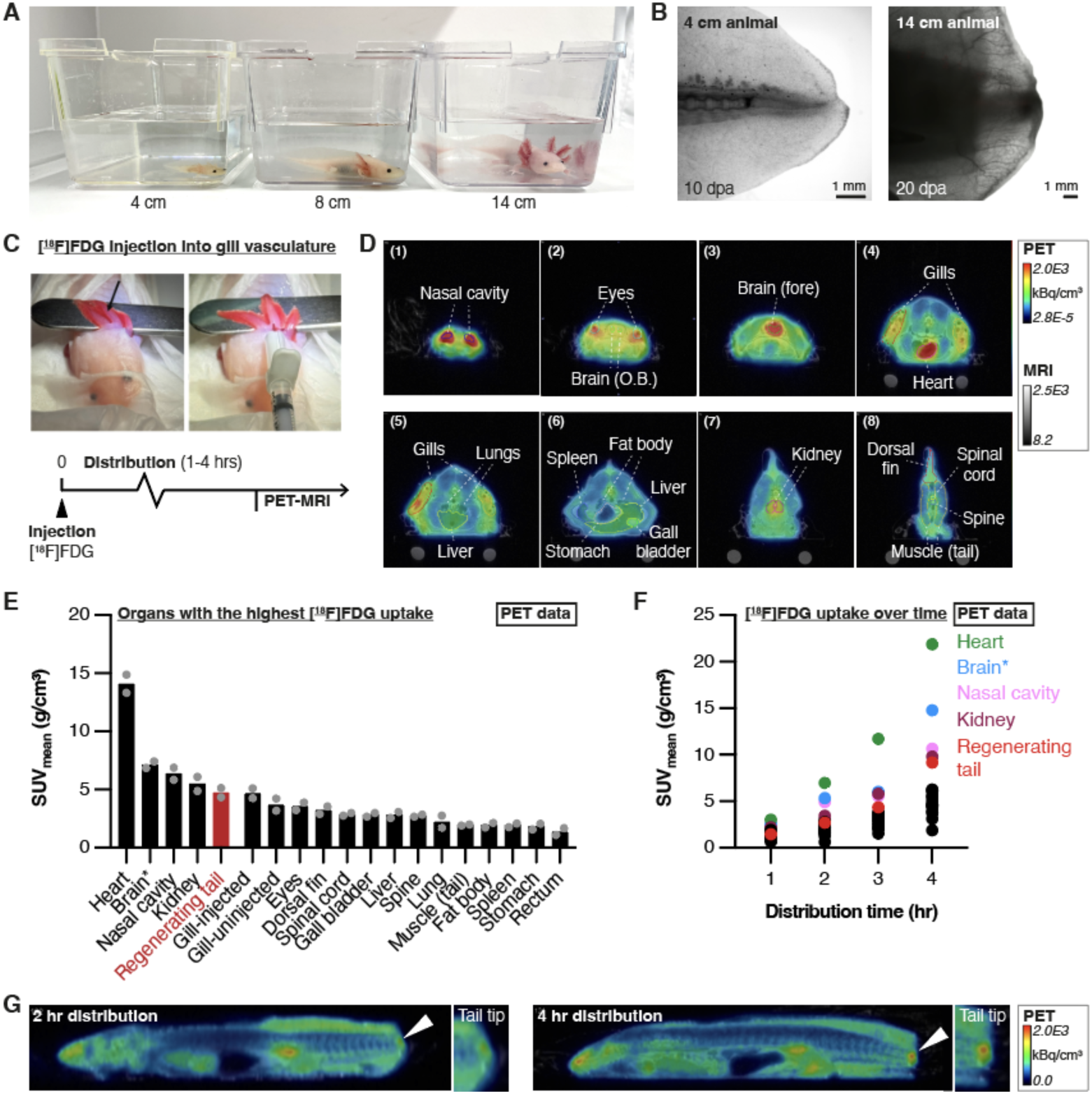
Application of PET-MRI to large, regenerating axolotls. (A) Axolotls of different sizes (animal lengths measured from snout to tail tip). (B) Internal tissues are visible with light microscopy in small axolotls (4 cm), but not in large axolotls (14 cm). Widefield images of live, regenerating axolotls. (C) Injection of [^18^F]FDG tracer into the blood vessels of the gill using an insulin syringe and 30 G needle. [^18^F]FDG was allowed to distribute systemically for between 1 and 4 hours in pilot experiments. (D) Axial (cross-section) reconstruction showing [^18^F]FDG accumulation in selected organs of interest. Organs were segmented based on MRI data (not shown) and panels are ordered from snout to tail. Red colour indicates higher signal and blue colour indicates lower signal. O.B.: olfactory bulb. Fore: forebrain. Representative axolotl 4 hours after [^18^F]FDG injection. (E) Mean standard uptake values (SUV_mean_) in selected organs 3 hours after injection of [^18^F]FDG, as measured using PET-MRI. Arranged in decreasing order. The regenerating tail tip is highlighted in red. *n* = 2 for pilot experiments. Brain*: forebrain and midbrain only. (F) SUV_mean_ in selected organs after [^18^F]FDG distribution for 1 – 4 hours. Organs with the highest uptake values are labelled. The same animal was measured at 1 and 3 hours, or 2 and 4 hours. (G) Representative PET images of the same axolotl imaged 2 and 4 hours after [^18^F]FDG injection. Arrowheads indicate signal in the regenerating tail tip, which is more obvious at 4 hours than 2 hours. Insets are magnifications of the tail tip. Sagittal view, head pointing left.

To overcome this limitation of optical imaging, we used FDG-PET/MRI to investigate glucose metabolism in larger animals. Axolotls are well-suited to PET/MRI, as they can be kept immobilised and hydrated by wrapping in anaesthetic-soaked tissue during imaging ^21^. However, as ectothermic amphibians, axolotls require adaptations to radiotracer administration and distribution time compared with standard rodent protocols ^22^. We therefore performed pilot experiments in a small number of male axolotls to minimise potential sex-related variation. After establishing suitable imaging conditions, we conducted regeneration experiments in both male and female axolotls. Details of all axolotls used for metabolic imaging are provided in **Table 1**.

**Table 1.** Details of axolotls depicted in figures for PET/MRI and PET/CT experiments. The lengths of injured axolotls are recorded as: original length – amputated length. NR: not recorded.

| Figure | Axolotl ID | Sex | Weight (g) | Length (cm) | Injury type | [<sup>18</sup>F]FDG distribution |
| --- | --- | --- | --- | --- | --- | --- |
| **2D** | A026 | M | 20.9 | 14.2 | Uninjured | 4 hr |
| **2E** | A013 | M | 19.4 | 14 - 4 | 14 dpa | 3 hr |
|  | A016 | M | 20.1 | 14 - 4 | 14 dpa | 3 hr |
| **2F** | A012 | M | 21.0 | 14 - 4 | 14 dpa | 2 hr / 4 hr |
|  | A015 | M | 18.1 | 14 - 4 | 14 dpa | 1 hr / 3 hr |
| **2G** | A012 | M | 21.0 | 14 - 4 | 14 dpa | 2 hr / 4 hr |
| **3D** | A027 | M | 20.2 | 13.9 - 2 | 7 dpa | 4 hr |
|  | A039 | F | 20.9 | 13.4 - 2 | 28 dpa | 4 hr |
|  | A022 | F | 18.5 | 13.7 - 4 | 7 dpa | 4 hr |
|  | A035 | M | 19.1 | 13.1 - 4 | 28 dpa | 4 hr |
| **3E** | *Full experimental series (see below)* |  |  |  |  |  |
| **4A** | *Full experimental series (see below)* |  |  |  |  |  |
| **4B** | *Full experimental series (see below)* |  |  |  |  |  |
| **4C** | *Full experimental series (see below)* |  |  |  |  |  |
| **S1B** | A002 | NR | 20.0 | 15 | Uninjured | 90 mins water bath |
|  | A003 | NR | 18.0 | 15 - 4 | 11 dpa | 90 mins water bath |
| **S2A** | A026 | M | 20.9 | 14.2 | Uninjured | 4 hr |
| **S2B** | A030 | F | 18.6 | 13.4 - 2 | 28 dpa | 4 hr |
|  | A011 | M | 37.0 | 15 | Uninjured | 3 hr |
| **S2C** | A011 | M | 37.0 | 15 | Uninjured | 3 hr |
|  | A014 | F | 21.3 | 14 | Uninjured | 3 hr |
| **S2D** | A013 | M | 19.4 | 14 - 4 | 14 dpa | 3 hr |
|  | A016 | M | 20.1 | 14 - 4 | 14 dpa | 3 hr |
| **S2E** | A012 | M | 21.0 | 14 - 4 | 14 dpa | 2 hr / 4 hr |
|  | A013 | M | 19.4 | 14 - 4 | 14 dpa | 3 hr |
|  | A015 | M | 18.1 | 14 - 4 | 14 dpa | 1 hr / 3 hr |
|  | A016 | M | 20.1 | 14 - 4 | 14 dpa | 3 hr |
| **S2F** | A015 | M | 18.1 | 14 - 4 | 14 dpa | 1 hr / 3 hr |
| **S3A** | A030 | F | 18.6 | 13.4 - 2 | 26 dpa | None |
|  | A031 | F | 18.5 | 12.2 - 4 | 26 dpa | None |
|  | A032 | M | 18.4 | 13.7 - 4 | 26 dpa | None |
|  | A033 | F | 20.2 | 13.2 - 4 | 25 dpa | None |
|  | A034 | F | 20.7 | 12.5 - 2 | 25 dpa | None |
|  | A035 | M | 19.1 | 13.1 - 4 | 25 dpa | None |
|  | A036 | M | 23.4 | 14.3 - 2 | 26 dpa | None |
|  | A037 | F | 19.4 | 13.3 - 4 | 26 dpa | None |
|  | BU1 | M | 20.7 | 13.2 - 2 | 26 dpa | None |
|  | A039 | F | 20.9 | 13.4 - 2 | 25 dpa | None |
|  | A040 | M | 21.3 | 13.5 - 2 | 25 dpa | None |
|  | BU2 | M | 22.3 | 13.5 - 4 | 25 dpa | None |
| **S3B** | *Full experimental series (see below)* |  |  |  |  |  |
| **S3C** | A042 | M | 23.7 | 14 | Uninjured | None |
|  | A043 | M | 24.6 | 14.2 | Uninjured | None |
|  | A044 | M | 25.8 | 14.2 | Uninjured | None |
|  | A045 | M | 24.2 | 13.9 | Uninjured | None |
| **S3D** | *Full experimental series (see below)* |  |  |  |  |  |
| **S4A** | *Full experimental series (see below)* |  |  |  |  |  |
| **S4B** | *Full experimental series (see below)* |  |  |  |  |  |
| **S4C** | *Full experimental series (see below)* |  |  |  |  |  |
| **Full experimental series** |  |  |  |  |  |  |
| Uninjured | A026 | M | 20.9 | 14.2 | Uninjured | 4 hr |
|  | A029 | M | 19.8 | 13.9 | Uninjured | 4 hr |
|  | A038 | F | 25.0 | 14.1 | Uninjured | 4 hr |
|  | A041 | M | 23.6 | 14 | Uninjured | 4 hr |
| Small injury, 7 dpa | A018 | F | 19.1 | 13.8 - 2 | 7 dpa | 4 hr |
|  | A019 | M | 17.7 | 14.1 - 2 | 7 dpa | 4 hr |
|  | A021 | M | 19.8 | 14 - 2 | 7 dpa | 4 hr |
|  | A024 | F | 18.2 | 13.4 - 2 | 7 dpa | 4 hr |
|  | A027 | M | 20.2 | 13.9 - 2 | 7 dpa | 4 hr |
| Small injury, 28 dpa | A030 | F | 18.6 | 13.4 - 2 | 28 dpa | 4 hr |
|  | A034 | F | 20.7 | 12.5 - 2 | 28 dpa | 4 hr |
|  | A036 | M | 23.5 | 14.3 - 2 | 28 dpa | 4 hr |
|  | A039 | F | 20.9 | 13.4 - 2 | 28 dpa | 4 hr |
|  | A040 | M | 21.3 | 13.5 - 2 | 28 dpa | 4 hr |
| Large injury, 7 dpa | A020 | M | 18.8 | 13.4 - 4 | 7 dpa | 4 hr |
|  | A022 | F | 18.5 | 13.7 - 4 | 7 dpa | 4 hr |
|  | A023 | M | 18.1 | 13.5 - 4 | 7 dpa | 4 hr |
|  | A025 | F | 17.4 | 13.4 - 4 | 7 dpa | 4 hr |
|  | A028 | F | 19.0 | 13.7 - 4 | 7 dpa | 4 hr |
| Large injury, 28 dpa | A031 | F | 18.5 | 12.2 - 4 | 28 dpa | 4 hr |
|  | A032 | M | 18.4 | 13.7 - 4 | 28 dpa | 4 hr |
|  | A033 | F | 20.2 | 13.2 - 4 | 28 dpa | 4 hr |
|  | A035 | M | 19.1 | 13.1 - 4 | 28 dpa | 4 hr |
|  | A037 | F | 19.4 | 13.3 - 4 | 28 dpa | 4 hr |

### PET/MRI reveals glucose uptake in large, regenerating axolotls

We first attempted to deliver FDG by bathing, an established method for delivering chemicals into axolotls ^26,27^. However, bathing was ineffective at delivering the radiotracer systemically within a reasonable timeframe, resulting in overall low accumulation throughout the body, with only slight enhancement in the heart and nearby tissues (**Figures S1A-B**). By contrast, we found that directly injecting FDG into the vasculature resulted in a well-distributed signal throughout the body. We injected into the blood vessels of the gills (**Figure 2C**), which are distant from the tail and minimise the risk that radiotracer trapped at the injection site interferes with measurements. We performed PET imaging at 1-4 hours after injection and simultaneously acquired anatomical MRI data to quantify tracer uptake in different organs (**Figures 2D, S2A**, and **Videos 1-2**). Whole-body imaging of 14 cm axolotls within a single field of view enabled accurate reconstruction of anatomy, with a PET voxel size of 500 μm and an MRI voxel size of 125 μm. MRI further resolved internal structures not central to this study, including the gonads (**Figure S2B**), enabling sex determination, and the lungs, which we observed in either inflated or collapsed states (**Figure S2C**).

We quantified standardized uptake values (SUV) for each organ. The organs with the highest FDG uptake were heart (14.09 g/cm^3^), “brain” (forebrain plus midbrain in this study; 7.13 g/cm^3^), nasal cavities (6.37 g/cm^3^), and kidneys (5.49 g/cm^3^) (**Figure 2E**). These sites are similar to those found in mammalian models. Excitingly, the regenerating tail tip was the tissue with the next highest uptake (4.25 g/cm^3^) (**Figure 2E**), indicating that the metabolic activation we had observed in small axolotls is also present in large axolotls and detectable using PET/MRI. These results were validated using a complementary *ex vivo* assay in which individual dissected organs were analysed by gamma counting (**Figure S2D**). A minor degree of tracer loss into the surrounding water was observed, likely due to leakage or excretion (**Figure S2E**).

To define the optimal distribution time for FDG, we performed PET/MRI on the same axolotls at 1 and 3 hours, or 2 and 4 hours after injection. We found that FDG uptake was highest after 4 hours in all measured organs (**Figure 2F**). The mean increase in uptake between 1 and 3 hours was 1.33 ± 0.46-fold across all measured organs, and 1.56 ± 0.36-fold between 2 and 4 hours. When analysing the PET images, local accumulation of FDG in the amputated tail tip was clearly detectable after 4 hours, but less so at 1, 2 or 3 hours (**Figures 2G, S2F**). A longer distribution time leads to greater tracer loss from radioactive decay (^18^F half-life: 109.8 mins), but this was offset by improved accumulation of tracer in target organs.

In summary, we successfully detected glucose uptake in regenerating axolotls *in vivo* using PET/MRI. Based on these pilots, we delivered FDG via the gill vasculature and allowed a 4-hour distribution time in subsequent experiments.

### Larger injuries induce higher glucose uptake than smaller injuries

Having established appropriate conditions, we used PET/MRI to test whether injury size affects the metabolic response during regeneration. Larger injuries were induced by more proximal tail amputations, while smaller injuries were generated by more distal amputations, allowing injury size to be varied over several centimetres. Interestingly, newts regenerate small and large tail injuries within a similar overall timeframe, despite substantial differences in the amount of tissue that must be replaced ^28^. This means that larger injuries regenerate relatively faster. Previous work has implicated differential cell cycle control in tail regeneration rate ^16^, although whether metabolic differences also contribute remains unclear.

In axolotls, myotomes (muscle segments) are regenerated within a similar timeframe following small or large tail injuries ^16^. We found a similar pattern in the axolotl spinal cord, which extended more rapidly after larger injuries than smaller ones, eventually recovering the lost length within ∼36 days in both conditions (**Figures 3A-C**). Faster tail regeneration was also seen after large injuries in the 14 cm axolotls used for PET/MRI experiments, with no significant difference between male and female animals (**Figure S3A**).

**Figure 3.**
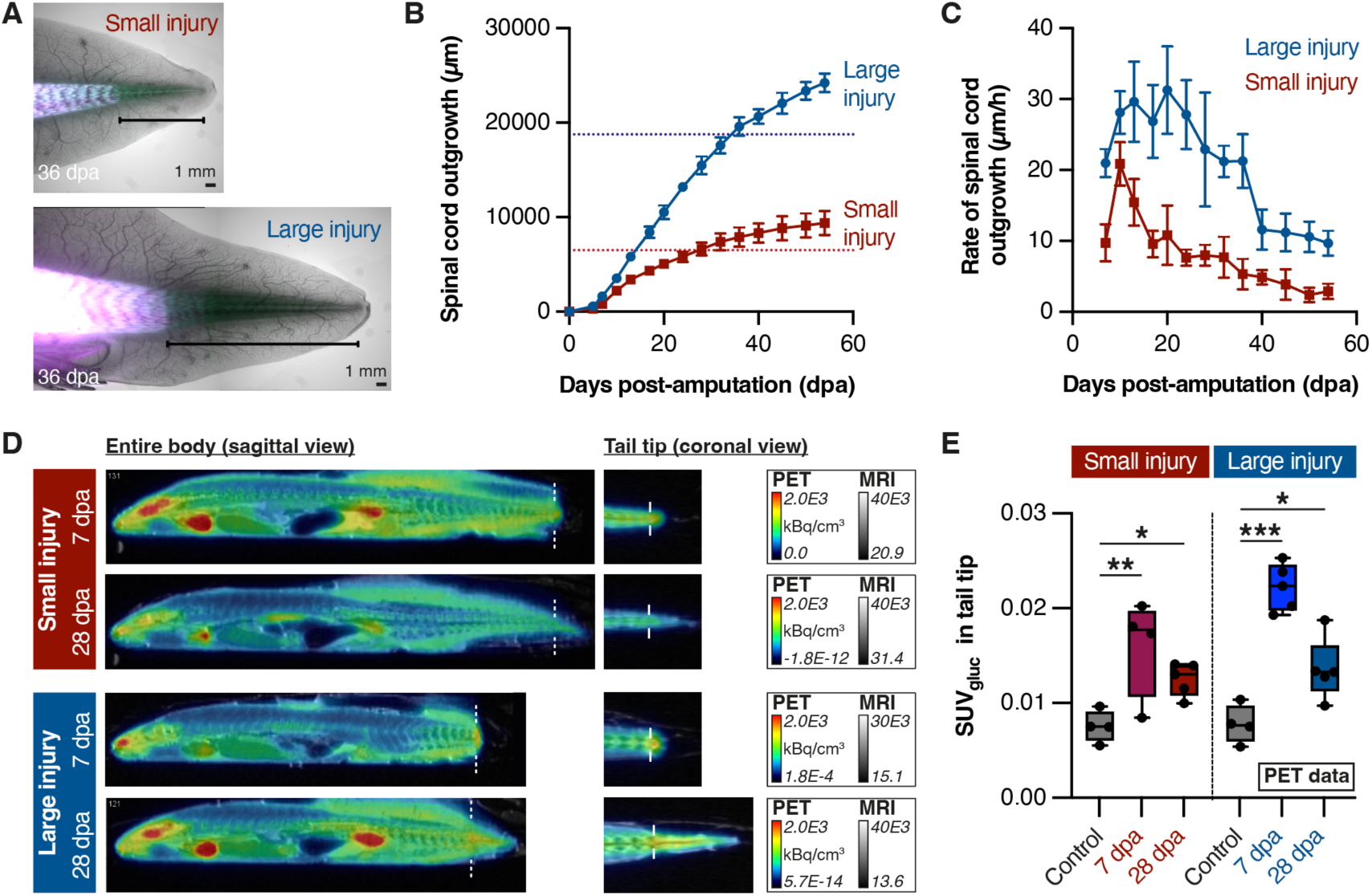
Injury size determines local and systemic glucose uptake during regeneration. (A) Widefield images depicting the amount of tail regenerated 36 days after a small injury (distal cut) or a large injury (proximal cut). The bracket indicates the regenerated region. The fluorescent reporter is AxFUCCI (Cura-Costa, Otsuki et al. 2021) and is used here solely as a visual aid to indicate the amputation plane. (B) Quantification of spinal cord outgrowth beyond the amputation plane. Dotted lines indicate the original length of the spinal cord prior to amputation. *n* = 4 animals (large injury) or 6 animals (small injury). (C) Rate of spinal cord outgrowth after large or small injury, calculated using the data in (B). (D) Representative PET-MRI images of axolotls in injury cohorts. Dotted lines indicate amputation plane. (E) Quantification of [^18^F]FDG uptake at the regenerating tail tip after small or large injury compared to the equivalent region of uninjured tails in control animals. SUV was corrected for blood glucose concentration, resulting in SUV_gluc_ (g/cm³ˑmg/dL). One-way ANOVA with Šidák’s multiple comparisons within each injury size. From left to right: **: *p* = 1.06×10^-3^, *: *p* = 3.91×10^-2^, ***: *p* = 6.68×10^-7^, *: *p* = 1.63×10^-2^.

For main series PET/MRI experiments, we removed 2 cm (small injury) or 4 cm (large injury) of tail tissue and prepared replicate cohorts to analyse at 7 dpa and 28 dpa (*n* = 5 animals per cohort, except small injury 7 dpa, for which *n* = 4). 7 dpa represents an early time point during blastema formation, while 28 dpa reflects a timepoint at which ∼1 cm of tail has regenerated, with re-differentiation of some tissue, but incomplete regeneration. As controls, we prepared a cohort of uninjured sibling axolotls (*n* = 4 animals).

As expected, we observed FDG accumulation at regenerating tail tips in injury cohorts, but not in controls (**Figure 3D**). We quantified uptake in the blastema. In these experiments, we measured blood glucose and corrected the SUV extracted from the PET images using these values (SUV_gluc_), as endogenous blood glucose competes with FDG, which can result in underestimated uptake values. We found that regenerating tail blastemas had significantly higher SUV_gluc_ than controls, regardless of injury size or timepoint (**Figure 3E**). Interestingly, large injuries triggered a higher SUV_gluc_ at 7 dpa than small injuries (mean SUV_gluc_ 0.022 (large) *vs* 0.016 (small), *p* = 5.09 x 10^-3^, one-way ANOVA with Šidák’s multiple comparisons) (**Figure 3E**). By 28 dpa, SUV_gluc_ in large injuries decreased to the same level as small injuries, although both remained elevated compared to controls (mean SUV_gluc_ 0.014 (large) and 0.013 (small) compared to 0.008 (both controls), *p* = 0.02 and 0.04 respectively, one-way ANOVA with Šidák’s multiple comparisons). There was no statistical difference in this response between male and female axolotls (mixed-effects analysis). Thus, large injuries trigger higher glucose uptake at the tail tip during early regeneration, correlating with faster regeneration speed. At later stages, FDG uptake per tissue volume is similar in large and small injuries.

### Elevated blood glucose levels do not affect the conclusions

During these analyses, we found that the blood glucose concentrations we measured were higher than previously reported values for axolotls (**Figure S3B**) ^29^. We hypothesised that the relatively long anaesthesia times in our protocol might be responsible for this, given similar findings in rodent models ^30,31^. Indeed, we found in separate experiments that a longer exposure to benzocaine correlated with higher blood glucose in axolotls (**Figure S3C**). We cannot rule out additional contributors, including noise/handling during PET/MRI, as well as repeated needle pricks for FDG administration and blood glucose measurement. Nevertheless, we performed all experiments with minimal variation, both in overall exposure to anaesthesia (97 mins ± 9 mins) and in total handling time from start to end of the experiment (5 hrs 5 mins ± 10 mins, *n* = 24 axolotls). Most importantly, blood glucose levels were not responsible for our conclusions. We confirmed that the relative differences that we had found between the experimental groups were preserved regardless of correction for blood glucose (**Figure 3E, S3D**).

### Systemic changes in glucose uptake after injury

It has recently been reported that injury to the axolotl limb induces body-wide (systemic) changes in cell proliferation and physiology in organs beyond the regenerating appendage ^32,33^. Our PET/MRI approach was well suited to detect a similar response in metabolism following tail injury. We therefore quantified SUV_gluc_ values across multiple organs using PET/MRI and validated these measurements using *ex vivo* gamma counter analysis as an independent method. In particular, we were interested in whether systemic responses would differ between small and large injuries.

As a pre-requisite for this analysis, we confirmed that FDG was well-distributed in the body, without trapping at the gill injection site. We found no significant difference in SUV_gluc_ values between injected and non-injected gills, indicating efficient tracer circulation (ratios between SUV_gluc_ in non-injected/injected gills were 0.94, 0.99, 0.86, 0.99 and 0.95 for uninjured, small 7 dpa, small 28 dpa, large 7 dpa and large 28 dpa cohorts respectively) (**Figures 4A, S4A**). Next, we analysed the other organs in the body. Strikingly, we found that mean FDG uptake across all quantified organs increased significantly in regenerating animals compared to uninjured controls (**Figures 4B, S4B**). Further comparisons among individual organs revealed increased FDG uptake at 7 dpa in both the small- and large-injury groups. However, the increase was more pronounced after a large injury: significant increases were observed in the heart, brain, nasal cavities, kidneys, dorsal fin, spinal cord, gall bladder and spine (**Figure 4C**). This rather non-specific pattern suggests a systemic response that extends far beyond the injured tail. Such a strong response is not observed in small injuries, suggesting a link between injury size and metabolic response. By 28 dpa, almost all organs had returned to steady-state levels in both injury cohorts (**Figure 4C**). Importantly, gamma counter measurements were consistent with the PET data, showing similar uptake patterns (**Figure S4C**). Altogether, our results indicate that injury size affects local and systemic glucose allocation during axolotl regeneration.

**Figure 4.**
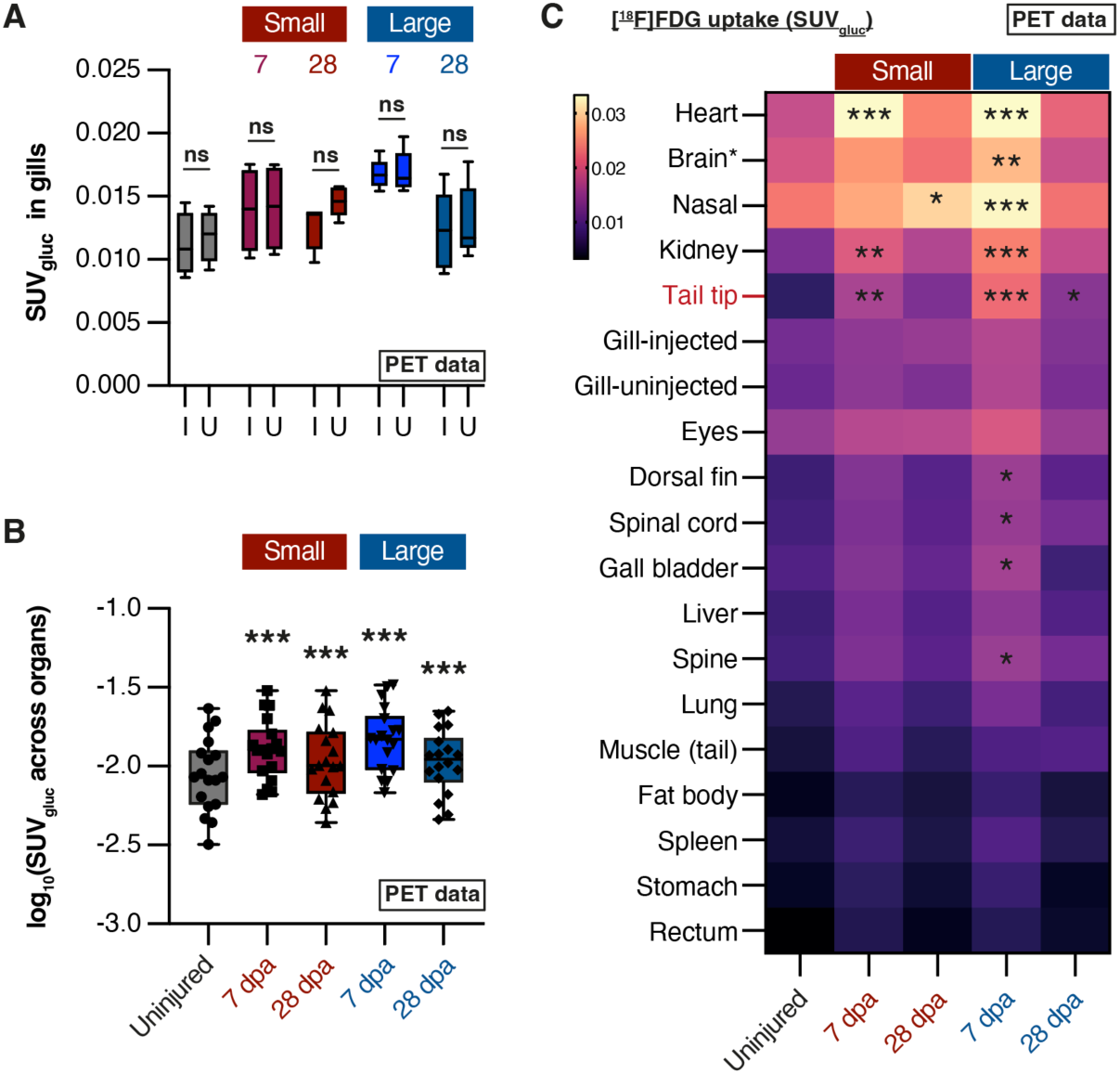
Tail injury triggers body-wide changes in glucose metabolism. (A) Paired analysis of [^18^F]FDG uptake in injected gills (I) and uninjected (U) gills in each experimental group, measured from PET data. No significant difference (ns) was seen in any group. Multiple paired *t*-tests, *p* > 0.05. *n* = 4 animals (control, small 7 dpa) or *n* = 5 animals (all other conditions). (B) Comparison of average SUV_gluc_ values (g/cm³ˑmg/dL) across experimental groups (PET data). Each dot represents a different organ, and represents the mean value from all animals in that experimental group. The regenerating tail tip was not included in these data. Data were log-transformed to allow statistical comparison by 2-way ANOVA with Dunnett’s multiple comparisons. From left to right, ***: *p* < 1×10^-12^, ***: *p* = 2.45×10^-8^, ***: *p* < 1×10^-12^, ***: *p* = 3.11×10^-7^. (C) Heatmap comparing SUV_gluc_ (g/cm³ˑmg/dL) in individual organs in each experimental group (PET data). Displayed heatmap value is the mean SUV_gluc_ of all animals in that experimental group. Yellow colours indicate higher values, while purple colours indicate lower values. Statistical significance compared to uninjured controls was determined by 2-way ANOVA with Dunnett’s multiple comparisons. *: *p* < 0.05, **: *p* < 0.01, ***: *p* < 1×10^-3^.

## Discussion

Recent studies have highlighted a conserved and essential role for glucose metabolism in regenerating diverse tissues in zebrafish and *Xenopus* species ^1–6^. An important question has been whether similar metabolic requirements apply to the regeneration of larger vertebrate tissues. Our results suggest that this is the case. Blocking glucose metabolism using 2-DG inhibited tail regeneration in axolotls. 2-DG has been widely used to demonstrate the importance of glucose metabolism for regeneration in other models ^1–6^ but it should be noted that it can broadly affect glycolysis and other glucose-dependent processes. Nevertheless, our results indicate that after tail loss, cells in proximity to the injury increase their glucose uptake and use this to induce regeneration. Recent studies in mouse embryos and in embryo models have shown that glycolytic metabolism can directly control cell fate decisions and signalling by regulating pathways such as Nodal and Wnt signalling ^34,35^. These roles are separable from the canonical functions of glycolysis in cellular energy. An intriguing possibility is that glycolytic metabolism similarly regulates signalling pathways during tissue regeneration.

Whole-body metabolic imaging revealed that injury size controls glucose allocation at both local and systemic levels. Larger injuries induce higher glucose uptake in the tail blastema than smaller injuries, consistent with faster regeneration, and also promote glucose uptake in distant organs. We do not rule out the possibility that the systemic effect reflects a generic physiological stress response induced by tissue loss. However, a recent study showed that limb amputation triggers a systemic stem cell activation response coordinated by adrenergic and mTOR signalling ^32,33^. Additionally, it has been reported that cryoinjury to the axolotl heart induces systemic upregulation of oxygen consumption and reduction in plasma lactate ^13^. Thus, our findings support emerging evidence that local injury elicits body-wide responses, some of which are relevant for regeneration. In the context of tail regeneration, these changes are present at 7 dpa but less prominent at 28 dpa. We have not determined whether these changes are instructive or supportive of regeneration, although immunostaining for pRPS6 suggests that metabolic changes occur in most blastema cells regardless of cell type or lineage. It will be important to identify how the glucose is used by axolotl cells, such as in the pentose phosphate pathway or hexosamine biosynthesis.

Previous studies of salamander metabolism are scarce and have focused on limb and heart regeneration ^8–13^. Global metabolic profiling suggested that limb blastema cells increase aerobic glycolysis during early regeneration, while shifting towards the TCA cycle later during differentiation ^10^. However, these correlations were not tested functionally, and no similar profiling has been performed on regenerating salamander tails. Although we did not perform unbiased profiling of metabolites, we spatially profiled glucose allocation in the body of an entire, regenerating vertebrate. Due to the spatial resolution limits of PET of around 1 mm³, a more detailed analysis would only be possible in dissected tissues *ex vivo*.

One limitation for both tail and limb regeneration is that injury size is intrinsically coupled to the proximal-distal axis of the organ (positional identity) if studied in one size of animal. A larger injury requires regeneration from a more proximal position (with a larger cross-sectional area), while a smaller injury regenerates from a more distal position (with a smaller cross-sectional area) ^16^. Tissue composition and the number of regenerative stem cells can vary along the proximal-distal axis. For example, the proximal tail (larger injuries) likely contains a higher proportion of muscle and other mesodermal tissue relative to epidermis when compared to the distal tail (small injuries). Although FDG-PET enables assessment of glucose metabolism in the whole body, the spatial resolution of PET imaging is insufficient to delineate glucose consumption at the level of individual cells or cell types. Thus, it will be necessary to use follow-up techniques to assess if the increase in glucose uptake in the tail after larger injuries reflects (i) the same number of progenitor cells, each taking up more glucose than after a smaller injury, (ii) a larger number of progenitor cells, each taking up the same amount of glucose as in a smaller injury or (iii) a combination of both.

We applied PET/MRI to a *bona fide* regenerating vertebrate and, in doing so, established a platform for investigating metabolic allocation during large-scale tissue regeneration. We showed that the metabolic response adapts to the extent of tissue loss and to changes beyond the injury site. The large size of axolotls (up to ∼ 30 cm) compared to other regenerative animal models makes them well-suited for PET/MRI as individual organs can be segmented and analysed. Extending this investigation to further PET tracers, such as lactate, amino acid or fatty acid radiotracers ^36–38^, will uncover the metabolic shifts occurring during regeneration. A major advantage of PET/MRI is that it is compatible with longitudinal studies of the same animal over time – potentially over the entire regeneration process. This will help in reconstructing the metabolic landscape for large-scale tissue regeneration.

## Methods

### Ethical oversight

Animal experiments were approved under licenses GZ:51072/2019/16, GZ:MA58-1432587-2022-12 and GZ:MA58-1516101-2023-21 by the Magistrate of Vienna (GMO office and MA58, Vienna, Austria) and GZ: 2023-0.185.563 by the Federal Ministry of Science, Research and Economy. PET, CT, MRI and gamma counter experiments were performed at the Preclinical Imaging Lab (PIL) of the Medical University of Vienna. All other experiments were performed at the Institute of Molecular Biotechnology (IMBA) and Research Institute of Molecular Pathology (IMP), Vienna BioCenter, Vienna.

### Axolotl husbandry

Axolotls (*Ambystoma mexicanum*) were spawned and raised in an in-house breeding colony at the Vienna BioCenter. Axolotl tanks were filled with Vienna tap water adjusted for conductivity and pH by the animal care team at the Vienna BioCenter. Axolotl sizes are reported in cm, measured from snout to tail tip, and weights are reported in grams. PET/MRI experiments were performed on approximately equal numbers of male and female animals, except pilot experiments that used only males. Details of all animals used in PET/MRI experiments, including animal sex, size and weight, are given in **Table 1**. Animal procedures were performed under anaesthesia in benzocaine (Merck, E-1501) diluted in tap water to a final concentration of 0.03% (PET/MRI) or 0.015% (other procedures). Animals received analgesia (butorphanol) treatment after tail amputation or intraperitoneal injection.

### Axolotl lines

Axolotls with different genetic backgrounds were used for the experiments. The *Sox2*:mCherry line was generated by ^24^ and originally referred to as *Sox2:Sox2*-ORFΔ-T2A-mCherry. Experiments in **Figures 3A-C** were performed on FUCCI axolotls ^14^. PET/MRI experiments were performed on sibling axolotls obtained from a cross between *Hand2*:EGFP and *Alx4*:mCherry parents ^39^. Other experiments were performed on control strain (*d/d*) axolotls.

### 2-DG experiments

4 cm *Sox2*:mCherry axolotls were injected intraperitoneally with 350 μg of either 2-DG (Sigma-Aldrich, D6134) or glucose (Sigma-Aldrich, G8270) (*n* = 5 axolotls per condition). The injection volume per animal was 20 μl, with 2-DG or glucose diluted in 70% PBS. Fast Green FCF dye was added to the injection mix to aid visualisation. 30 mins after injection, each animal had 0.5 cm of the distal tail tip amputated using a surgical scalpel. 2-DG/glucose injections were repeated every other day until 10 days post-amputation (10 dpa). Animals were imaged using an AXIOzoom V16 widefield microscope (Zeiss). Imaging was performed prior to injection on any given day.

### 2-NBDG experiments

6 cm *d/d* axolotls received a tail amputation 6 days before these experiments. Regenerating axolotls were anaesthetised, then injected intraperitoneally either with 1.5 mM 2-NBDG (Cayman Chemical, CAY-11046-5) or 70% PBS (control) (*n* = 3 axolotls per condition). The injection volume per animal was 20 μL. Animals were imaged using an AXIOzoom V16 widefield microscope (Zeiss) 2 hours after injection.

### Tissue harvesting, sectioning and immunostaining for pRPS6^Ser240/244^

14 cm *d/d* axolotls were used for these experiments (*n* = 4 axolotls per condition). 4 cm of the tail tip were amputated to induce regeneration. The regenerating blastema and adjacent tail stump were harvested at 7 dpa. For control samples, the equivalent tail region of uninjured sibling animals was harvested. Samples were fixed overnight at 6 °C in 4% paraformaldehyde, pH 7.4. After washing well in cold PBS, fixed samples were equilibrated at 6 °C for one night each with 20% sucrose and 30% sucrose. After this, samples were mounted in Tissue-Tek OCT compound (Sakura), frozen on dry ice and stored at −70 °C until sectioning. Cross sections of thickness 16 μm were prepared using a Cryostar NX70 cryostat and stored at −20 °C until staining. For immunostaining, slides were brought to room temperature before washing off the OCT with PBS. Slides were permeabilised with PBTx (PBS containing 0.2% Triton X-100), then blocked with 3% BSA in PBTx. Immunostaining was performed overnight at 6 °C with anti-pRPS6^Ser240/244^ antibody (CST4858) diluted 1:200 in PBTx + 0.3% BSA. The next day, slides were washed well with PBTx then incubated for 2 hours at room temperature with PBTx containing 1:500 anti-rabbit-Alexa 647 secondary antibody (Invitrogen) and DAPI. After final washes with PBTx, slides were mounted in Abberior Mount liquid antifade mounting media (Abberior). Images were acquired using a LSM980 AxioObserver inverted confocal microscope with ZEN software (Zeiss).

### Tail outgrowth speed measurements

7 cm FUCCI axolotls received a tail amputation removing either 0.7 cm of tail tip (24 myotomes post-cloaca, small injury, *n* = 6 axolotls) or 1.9 cm of tail tip (12 myotomes post-cloaca, large injury, *n* = 4 axolotls). Spinal cord outgrowth was measured from brightfield images acquired every few days during regeneration using an AXIOzoom V16 widefield microscope (Zeiss). The FUCCI fluorescent reporter was not used for quantifications – it is only displayed in the figure as a visual aid for the plane of amputation.

### Image analysis (widefield and confocal microscopy)

Microscope images were analysed using Fiji software version 2.3.0/1.53f ^40^.

### Pilot experiments with bath application of FDG (PET/Computed Tomography (CT))

15 cm axolotls (*n* = 3) were used, either uninjured or 11 days after a tail amputation removing 4 cm of tissue. Mean animal weight was 19.0 ± 1.0 g. Axolotls were incubated for 90 mins in a water bath containing 0.3 MBq/mL or 3 MBq/mL of FDG. The resulting total exposure was 50 MBq or 300 MBq respectively. Subsequently, axolotls were transferred to a new water bath containing clean water for 30 mins, then transferred to another tank containing 0.015% benzocaine anaesthesia for 20-30 mins. For FDG-PET/CT scanning, anaesthetised axolotls were removed from their tank, wrapped in a Kimtech towel soaked in fresh anaesthetic, and then placed onto the imaging cradle without further heating or anaesthesia.

### PET/CT experiments – technical specifications

PET/CT images were acquired using the Inveon MultiModality PET/CT scanner (Siemens Medical Solutions, Knoxville, TN, USA). The PET scanner has a field of view (FoV) of 100 mm (axial) and 127 mm (radial), a peak sensitivity of 6.7% and a spatial resolution (5 mm off-center FoV) of 1.8 mm, according to NEMA NU 4-2008 performance evaluation ^41^. CT scans (voltage 80 kVp, current 500 µA, exposure time 100 ms, 181 projections, scan time 3:16) were performed pre-injection of the tracer. CT images were reconstructed using Feldkamp cone beam and a binning 4, matrix size 512 x 512 x 512 resulting in voxel size 0.195127 mm³. CT images were calibrated to Hounsfield units (HU) using a scanner-specific calibration protocol. For PET scanning, emission data were recorded for 20 mins using an energy window of 350-60 keV, and a 3.4375 ns coincidence time window. PET images were reconstructed using an ordered subset expectation maximization (OSEM/MAP) algorithm with 18 iterations and 16 subsets and a matrix size of 256 x 256 x 159, resulting in a reconstructed voxel size of 0.0388 x 0.0388 x 0.0796 mm³. The default correction methods (normalization, CT-based attenuation, scatter, dead-time, decay correction) were applied to the PET data. Total scan time was around 24 mins (20:00 PET, 3:16 CT) as the PET and CT scans were performed sequentially.

### PET/MRI experiments - overview

For pilot experiments, 14 cm axolotls (*n* = 4) were used. All animals were male, received a large (4 cm) tail injury and were imaged at 14 dpa. Mean animal weight one day before imaging was 19.7 ± 1.1 g. FDG administration, PET/MRI and gamma counter measurements were carried out as similarly to the main series experiments below, with the following exceptions. Actual injected activity was 10.8 ± 4.2 MBq. Following the recommendation by Alstrup et al. ^22^, we initially scanned the animals at 3 hours post-injection (*n* = 2). However, to evaluate the time course of FDG distribution, 2 additional axolotls were scanned at both 1 and 3 hours (*n* = 1) or at both 2 and 4 hours (*n* = 1) post-injection.

For main series experiments, 14 cm axolotls were used (*n* = 24). Axolotls were allocated to the following experimental cohorts with approximately equal male-female balance: no injury (control); small injury (amputation of 2 cm of the tail tip); large injury (amputation of 4 cm of the tail tip). Axolotls were fasted without food for 2 days prior to PET/MRI experiments and were scanned at 14 dpa or 28 dpa. 3 animals were scanned per day, and replicates from each experimental condition were spread over multiple days to reduce bias in data acquisition. Mean animal weight was 19.8 ± 2.0 g.

The following description reports mean ± standard deviation (SD) values acquired from 24 animals in the main experimental series. Values from pilot experiments are not included here. Axolotls were anaesthetised individually in 400 mL of 0.03% benzocaine for 30 ± 6 mins. Anaesthesia depth was confirmed by checking whether the uprighting reflex was still present. Thereafter, the axolotl was taken out, wrapped in paper humidified with anaesthetic, and positioned under a microscope (Olympus SZ×10). The collapsed gills were positioned on a metal forceps handle for better stability. FDG was injected preferentially into the centre-left gill using an insulin syringe with 30G needle (20 - 30 μL containing 31.3 ± 13.5 MBq activity). The actual injected dose (16.2 ± 3.5 MBq) was estimated by immediately measuring the whole axolotl using a dose calibrator. Axolotls were returned to a tank containing 400 mL of fresh tap water and FDG was allowed to distribute for the desired time (4 hours for main experiments). To estimate loss of radioactivity by excretion and/or leakage, a 500 μL aliquot of tank water was sampled at the following hours after injection: 0.5, 1, 2, 3, 3.5. Axolotls were re-anaesthetised in the benzocaine bath starting 30 mins before PET-MRI scanning. For scanning, axolotls were removed from the bath, wrapped in a Kimtech towel soaked in anaesthetic then placed in prone position onto the imaging cradle, without heating, with the tail propped upright using polystyrene blocks. PET scanning was started 4 hours ± 1 min after FDG injection, simultaneous with the MRI. Total scan time was 30 ± 5 mins. After PET/MRI, the residual radioactivity in the whole animal was measured using the same dose calibrator (2.5 ± 0.5 MBq). A blood sample was taken from an uninjected gill to measure blood glucose concentration using a glucometer. Finally, the animal was euthanised by transecting the cervical spinal cord, individual organs were dissected out and radioactivity in each organ, plus the water aliquots, was measured using a gamma counter (see below). Total protocol time from first anaesthesia to sacrifice was 5 hours 5 mins ± 10 mins.

### PET/MRI experiments – technical specifications

PET/MRI was performed using a BIOSPEC® 94/30 USR MRI instrument (Bruker, Germany), equipped with a PET insert (PET Insert Si 198, Bruker, Germany). The PET insert has a field of view (FoV) of 150 mm (axial) and 80 mm (radial), a peak sensitivity of 10 % and a spatial resolution (5 mm off-center FoV) of 1.5 mm, according to NEMA NU4-2008 performance evaluation ^42^. For simultaneous PET and MR imaging, a quadrature birdcage transmit/receive RF coil with 86 mm inner diameter (Quad86, T20202V3) was used. All scans were acquired using the provided scanner software PARAVISION 360 V3.2. For PET scanning, an energy window of 30% (357.7-664.3 keV), and a 7 ns coincidence time window were applied. Emission data were recorded for 20 mins. Images were reconstructed using MLEM (18 iterations), a FoV of 90 x 90 x 150 mm³ and a matrix size of 180 x 180 x 300 yielding 0.5 mm isotropic voxel size. The default correction methods (normalization, scatter, dead-time, decay correction) were applied to the PET data. MR parameters for the different acquisitions were as follows: T2 RARE MR coronal: TE 24.24 ms, TR 8,000 ms, 2 averages, echo spacing 8.080, rare factor 8, FoV 135.294 x 30.00, slice thickness 0.5 mm, image size 256 x 256 yield a voxel size of 0.528 x 0.152 x 0.5 mm³. T2 RARE MR axial: TE 7.67 ms, TR 11,544 ms, 2 averages, echo spacing 7.667, rare factor 8, FoV 35.00 x 35.00, slice thickness 1.0 mm, image size 256 x 256 yield a voxel size of 0.137 x 0.137 x 1.0 mm³. T1 FLASH: TE 3.822 ms, TR 2,000 ms, 4 averages, flip angle 70°, FoV 35.00 x 35.00, slice thickness 1.0 mm, image size 280 x 72 yield a voxel size of 0.125 x 0.486 x 1.0 mm³.

### Image analysis (PET/MRI)

PET/MRI data were analysed using pmod Quantification Software (Bruker, version 4.4). PET and anatomical images were co-registered, then organs (volumes of interest, VOIs) were identified and manually delineated on consecutive axial or coronal planes. The person performing image analysis was blinded to the PET data while delineating VOIs in the MRI data. Details of how the VOIs were defined are given in **Table 2**. From the PET data, radioactivity concentrations were extracted for all defined VOIs and reported in standardized uptake values (SUVs; radioactivity concentration expressed in kBq/cm^3^ and corrected for injected activity and weight of the axolotl). In the main experiments, SUVs were corrected by multiplying by blood glucose concentration (mg/dL) to obtain SUV_gluc_ (g/cm³ˑmg/dL).

**Table 2.** Organ segmentation (delineation of VOIs) in MRI data. All segmentations were performed on MRI data, while blinded to PET data. All segmentations were performed on axially acquired data, except regenerating tail that was segmented in coronal data. Organs are arranged in alphabetical order.

| Organ / VOI | Criteria |
| --- | --- |
| Brain* | Forebrain and midbrain only. Areas for segmentation were defined based on the MRI brain atlas of Lazcano et al. 2021. |
| Dorsal fin | Excluding the outer skin layer. 10 planes of axial data (10 mm of tissue) were quantified posterior to the cloaca. |
| Eyes | Entire organ. |
| Fat body | Entire organ. Both fat bodies were quantified together. |
| Gall bladder | Entire organ. |
| Gill | The injected and uninjected gills were quantified separately. All fronds were segmented over 10 planes of axial data (10 mm of tissue). |
| Heart | Entire organ. |
| Kidney | Entire organ. Both kidneys were quantified together. |
| Liver | Entire organ. |
| Lung | Entire organ. Both lungs were quantified together. |
| Muscle (tail) | 10 planes of axial data (10 mm of tissue) were quantified posterior to the cloaca. Left and right muscle masses were quantified together. |
| Nasal cavity | External structures only. |
| Rectum | Entire organ. |
| Regenerating tail | All tail beyond the amputation plane, excluding the outer skin layer. The amputation plane was defined by the position of the severed spine. |
| Spinal cord | 10 planes of axial data (10 mm of tissue) were quantified posterior to the cloaca. |
| Spine | 10 planes of axial data (10 mm of tissue) were quantified posterior to the cloaca. |
| Spleen | Entire organ. |
| Stomach | Entire organ. |

### Tissue harvest for measurement with a gamma counter

After PET/MRI, axolotls were sacrificed by a cervical cut to the spinal cord. Individual organs were dissected out and collected separately to be measured using a gamma counter (Hidex). Dissections were performed with surgical scissors and forceps. **Table 3** lists the details of how each organ was dissected.

**Table 3.** Organ dissection for gamma counter analysis. All organs were manually dissected from axolotls that had undergone a PET/MRI scan. Organs are arranged in alphabetical order.

| Organ | Criteria |
| --- | --- |
| Blood (clot) | A sample of blood was taken, usually from a leakage during the dissection. |
| Brain* | Forebrain and midbrain as far as possible, although hindbrain contamination cannot be excluded. |
| Carcass | The remainder of the axolotl after removal of other organs. |
| Fat body + gonads | Entire organ, dissected together. |
| Gill | All three gill branches were harvested separately from the left and right sides of the animal. |
| Gut + Rectum | Entire organ, dissected together. Faeces were squeezed out of the rectum. |
| Heart | Entire organ. |
| Kidney | Entire organ. Both kidneys were quantified together. |
| Limb | Entire organ. All four limbs were quantified together. |
| Liver | Entire organ, with gall bladder removed. |
| Lung | Entire organ. Both lungs were quantified together. |
| Regenerating tail | The regenerating tail blastema was harvested. |
| Spine | A 1 cm piece of thoracic vertebral column was harvested, together with surrounding mesenchymal tissue. |
| Spleen | Entire organ. |
| Stomach | Entire organ. |
| Tail (control) | Regenerating animals: to avoid regenerating source tissue, 1 cm of tissue proximal to the amputation plane was dissected and discarded. Tissue further proximal than this was harvested. Uninjured controls: an equivalent region of the tail was harvested as in regenerating animals. |

### Gamma counter – technical specifications

Radioactivity in each organ was measured using a gamma counter and expressed in counts per minute (CPM) and converted to becquerels (Bq). These values were normalized to organ weight (g) to obtain radioactivity concentrations (Bq/g). To correct for inter-subject variability in administered radioactivity, the concentrations were further normalized to the total and corrected radioactivity applied to the axolotl, resulting in values expressed as %IA/g. Additionally, %IA/g_gluc_ was calculated by multiplying %IA/g by the blood glucose concentration of the respective animal (g/mL).

### Correlation between benzocaine anaesthesia and blood glucose levels

4 axolotls with a mean length of 14.1 ± 0.2 cm and a mean weight of 24.6 ± 0.9 g were used in this experiment. As this experiment was not carried out in parallel with the PET/MRI studies, the animals underwent a different treatment than in the main experiment. Each animal was anaesthetised in 0.03% benzocaine for 40 mins, then a blood sample was taken from one gill and measured using a glucometer (40 mins exposure). The animal was allowed to recover in water for 60 mins before returning to tank containing 0.03% benzocaine for a further 60 mins. A blood sample was taken for blood glucose measurement (100 mins total exposure). This step was repeated once more to obtain a blood glucose measurement after 160 mins exposure to 0.03% benzocaine. The animals were kept submerged at all times (no extraction and wrapping in a towel), and no imaging/scanning was performed.

### Data exclusions

The data from one animal in the small injury – 7 dpa cohort (A021) were acquired but excluded from statistical analyses as the amount of FDG injected in this animal was found to be 27.6% of the dose injected into the other animals of the same cohort. This exclusion was determined *a posteriori*.

### Additional experimental design

For PET/MRI experiments, animals were allocated *a priori* to yield approximately equal numbers of male and female individuals in each group (all axolotls are detailed in **Table 1**). Data were analysed blinded to sex. The operator performing the PET/MRI image analysis was aware of the animal’s allocated group (no blinding) but PET data were quantified within masks delineated solely on the MRI data (i.e. blinded to PET data). For all non-PET/MRI experiments, animals were allocated randomly and analyses were performed blinded to animal sex. Experimental details are reported in line with the ARRIVE Essential 10 guidelines ^43^.

### Statistical analysis and data representation

Measurements were taken from distinct animals without repeated imaging, except in the PET/MRI pilot experiments in which the same animal was imaged at 1 and 3 hours or 2 and 4 hours. Statistical analysis and graph plotting were performed using Prism software (GraphPad Prism, version 11.0.0). Data were tested for assumptions of normality and equality of variance to determine the appropriate statistical test to use, which is indicated in the legend of the respective Figure. Tests were adjusted for multiple comparisons. A threshold of *p* < 0.05 was used for statistical significance. Error bars indicate standard deviation. Mean values are reported ± standard deviation. Figures were assembled in Illustrator (Adobe).

## Additional information

## Acknowledgments

We thank the animal care team at the Vienna BioCenter for exceptional axolotl care: Viktoria Szilagyi, Andrea Lentz-Koblenc, Julia König, Dijana Bastian, Veronika Vojnicsek, Erika Kiligan, Elisabeth Zöllner, Daniela Pollak. We thank the Core facility laboratory animal breeding and husbandry at the Medical University of Vienna for the support with authorities to house axolotl during the experiments at the PIL. We thank the BioOptics facility at the Vienna BioCenter core facilities for expert microscopy support. We thank the following students of the Vienna BioCenter, Medical University of Vienna and the Hubrecht Institute for their help in preparing axolotls for PET/MRI: Willemijn Bout, Sarah Plattner, Viktoria Paller, Deniz Demirkesenler, Lisa Aichinger. We thank Johann Stanek for technical organisation at the PIL and also acknowledge the research platform medical imaging (RPMI).

## Open access

This research was funded in part by European Research Council (ERC) Advanced Grant 742046. For the purpose of open access, the authors have applied a CC BY public copyright licence to any Author Accepted Manuscript version arising from this submission.

## Author contributions

CK: Conceptualization, Data acquisition, Data analysis, Funding acquisition, Project supervision, Writing – original draft, Writing - editing.

CP: Conceptualization, Data acquisition, Data analysis, Funding acquisition, Project supervision, Writing – editing.

CV: Conceptualization, Data acquisition, Data analysis, Funding acquisition, Project supervision, Writing – editing.

LZ: Data acquisition, Writing - editing.

TW: Data acquisition, Data analysis.

JF: Data acquisition, Data analysis.

TH: Funding acquisition.

MH: Funding acquisition.

VW: Data acquisition, Data analysis.

EMT: Conceptualization, Data analysis, Funding acquisition, Writing – editing.

LO: Conceptualization, Data acquisition, Data analysis, Funding acquisition, Project supervision, Figure preparation, Writing – original draft, Writing – editing.

## Data availability

The experimental PET/MRI image data that support the findings of this study will be made available in the Preclinical Image DAtaset Repository

(PIDAR) https://pidar.hpc4ai.unito.it/Datasets/Index.

## Generative AI statement

ChatGPT (OpenAI) version 5.2 was used to lightly edit text for clarity after original writing by human authors.

## Video legends

**Video 1. Organ segmentation (MRI).**

Video depicting axolotl anatomy and segmentations of selected organs on axial MRI data. Video runs from snout to tail tip. The distance between adjacent sections is 1 mm. The same animal is depicted as in **Fig. 2D**.

**Video 2. [^18^F]FDG uptake in different organs (PET/MRI).**

Video depicting [^18^F]FDG uptake after 4 hours distribution in axial PET/MRI data. Video runs from snout to tail tip. The distance between adjacent sections is 1 mm. The same animal is depicted as in **Fig. 2D**.

## Supplementary Figures

**Figure S1.**
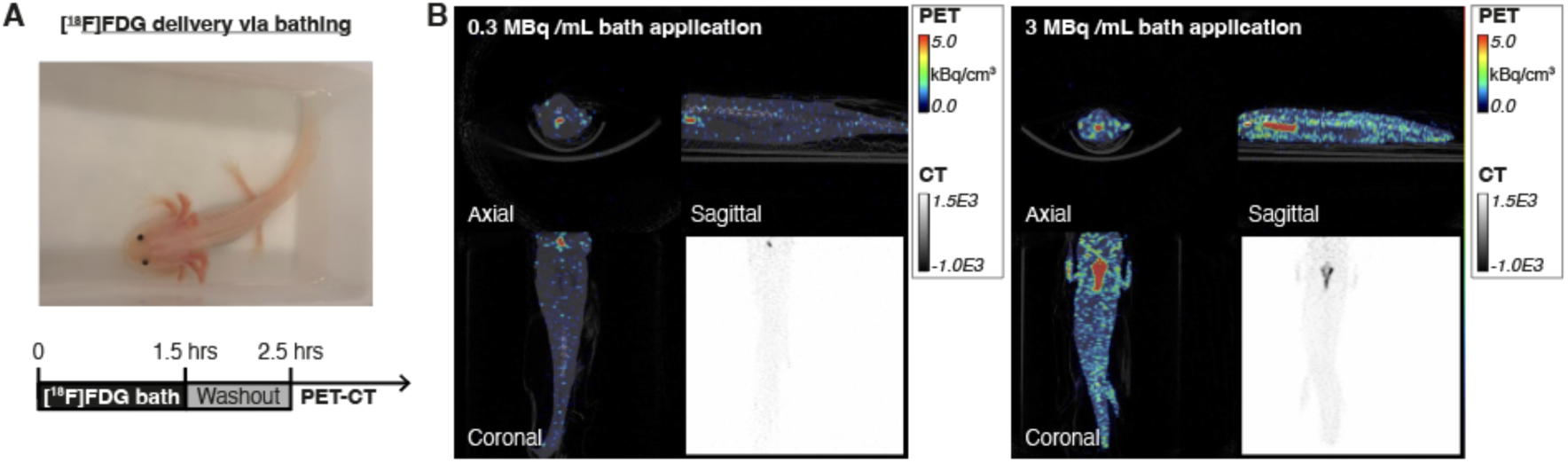
Bathing is an inefficient method to deliver [^18^F]FDG into axolotls in comparison to intravascular injection. (A) Image of a regenerating axolotl in a water bath containing [^18^F]FDG (above). Timeline summarising the experimental design (below). Only in this pilot experiment, PET imaging was paired with CT scanning to resolve skeletal structures (PET-CT). *n* = 3 animals. (B) PET-CT images depicting tracer distribution after bathing axolotls for 90 mins in a [^18^F]FDG-containing water bath with a radioactivity concentration of 0.3 MBq/mL (left) or 3 MBq/mL (right). The resulting total exposure was 50 MBq or 300 MBq respectively. Red indicates higher [^18^F]FDG activity; blue indicates lower activity.

**Figure S2.**
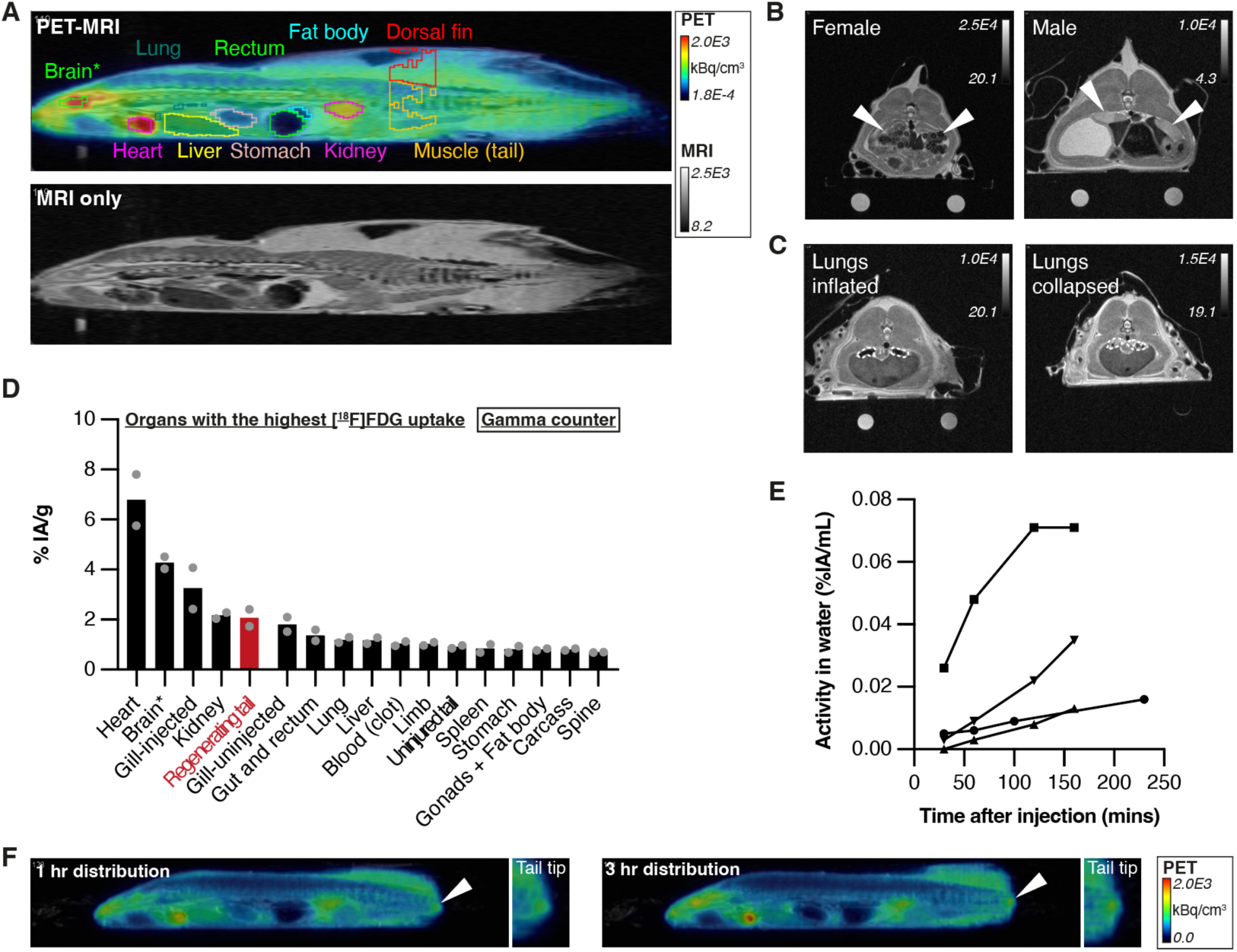
Establishment of PET-MRI conditions for regenerating axolotls. (A) A 14 cm axolotl 4 hours after injection with [^18^F]FDG, depicted in sagittal view. The tail tip is not visible in this image plane. Top image shows PET signal overlaid on MRI data, with segmented organs indicated. Bottom image shows MRI data only. (B) MRI images in axial view with reproductive structures arrowed. (C) MRI images in axial view with lungs outlined. (D) [^18^F]FDG dose in selected organs 3 hours after injection, as determined by gamma counter measurements of dissected organs. Units are % injected activity (IA)/g. Arranged in decreasing order. The regenerating tail tip is highlighted in red. *n* = 2 for pilot experiments. Brain*: forebrain and midbrain only. (E) Estimation of [^18^F]FDG leakage into the water bath (loss from the animal due to leakage or excretion). *n* = 4 animals. IA: injected activity. The volume of the water bath was 400 mL. (F) Representative PET images of the same axolotl imaged 1 and 3 hours after [^18^F]FDG injection. Arrowheads indicates signal in the regenerating tail tip. Sagittal view, head pointing left.

**Figure S3.**
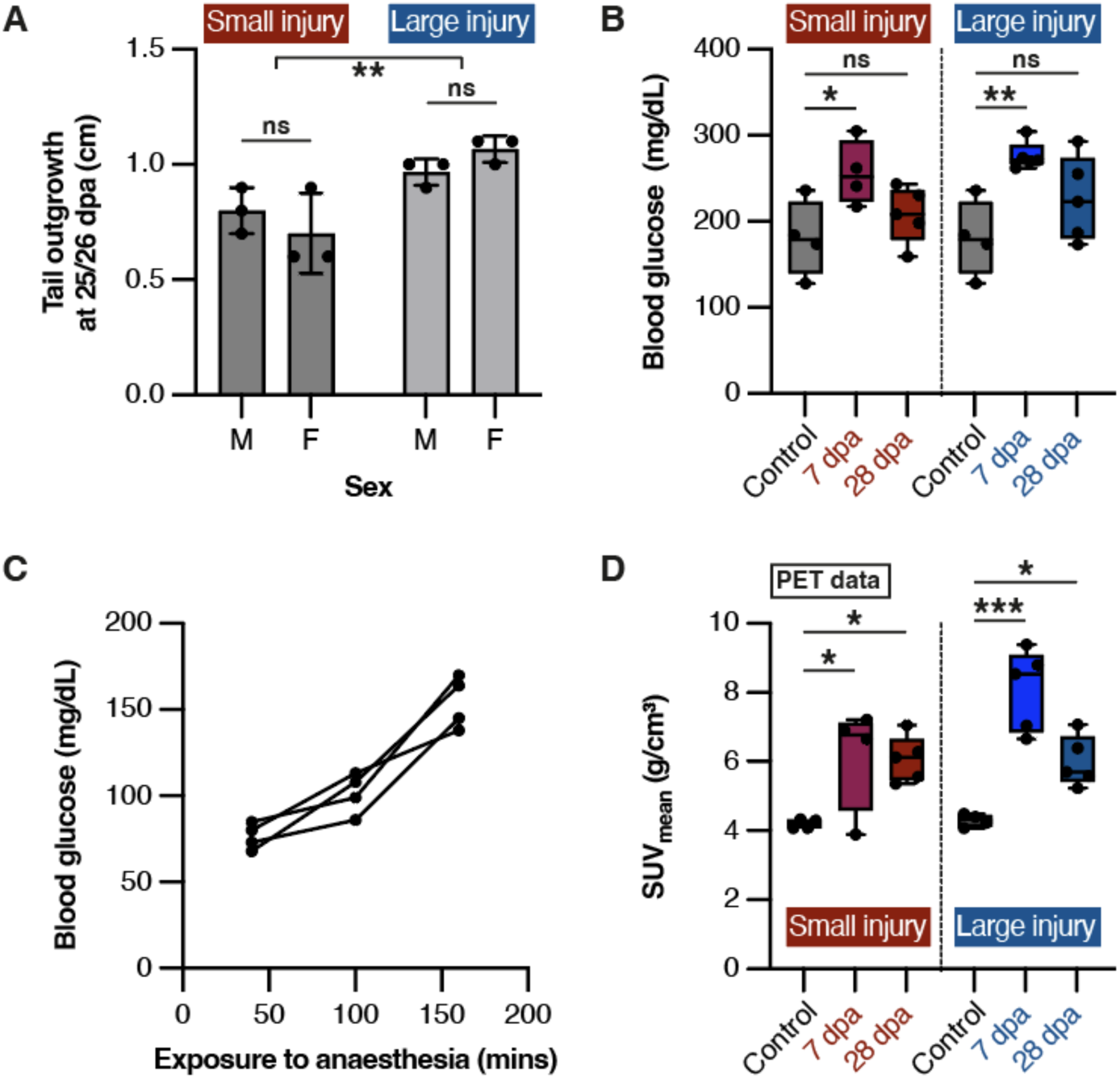
Characterisation of tail outgrowth and blood glucose effects. (A) Tail outgrowth at 25-26 days after small injury or large injury, separated by sex. Tails that received a large injury regenerated significantly more tissue than those that received a small injury (**: *p* = 2.70×10^-3^). No significant different (ns) was observed between male and female animals (*p* > 0.05). Two-way ANOVA, *n* = 3 animals per group (12 animals total). (B) Blood glucose levels in axolotls in each experimental cohort. One-way ANOVA followed by Šidák’s multiple comparisons test. *: *p* = 4.29×10^-2^, **: *p* = 5.06×10^-3^. (C) Blood glucose levels after increasing exposure to anaesthesia. Each line connects the datapoints from one animal. *n* = 4 animals. (D) Quantification of [^18^F]FDG uptake at the regenerating tail tip after small or large injury compared to the equivalent region of uninjured tails in control animals. SUV was not corrected for blood glucose concentration. One-way ANOVA, followed by a Holm-Šidák’s multiple comparisons test. From left to right: *p* = 2.08×10^-2^, *p* = 1.97×10^-2^, *p* = 1.10×10^-5^, *p* = 3.82×10^-2^.

**Figure S4.**
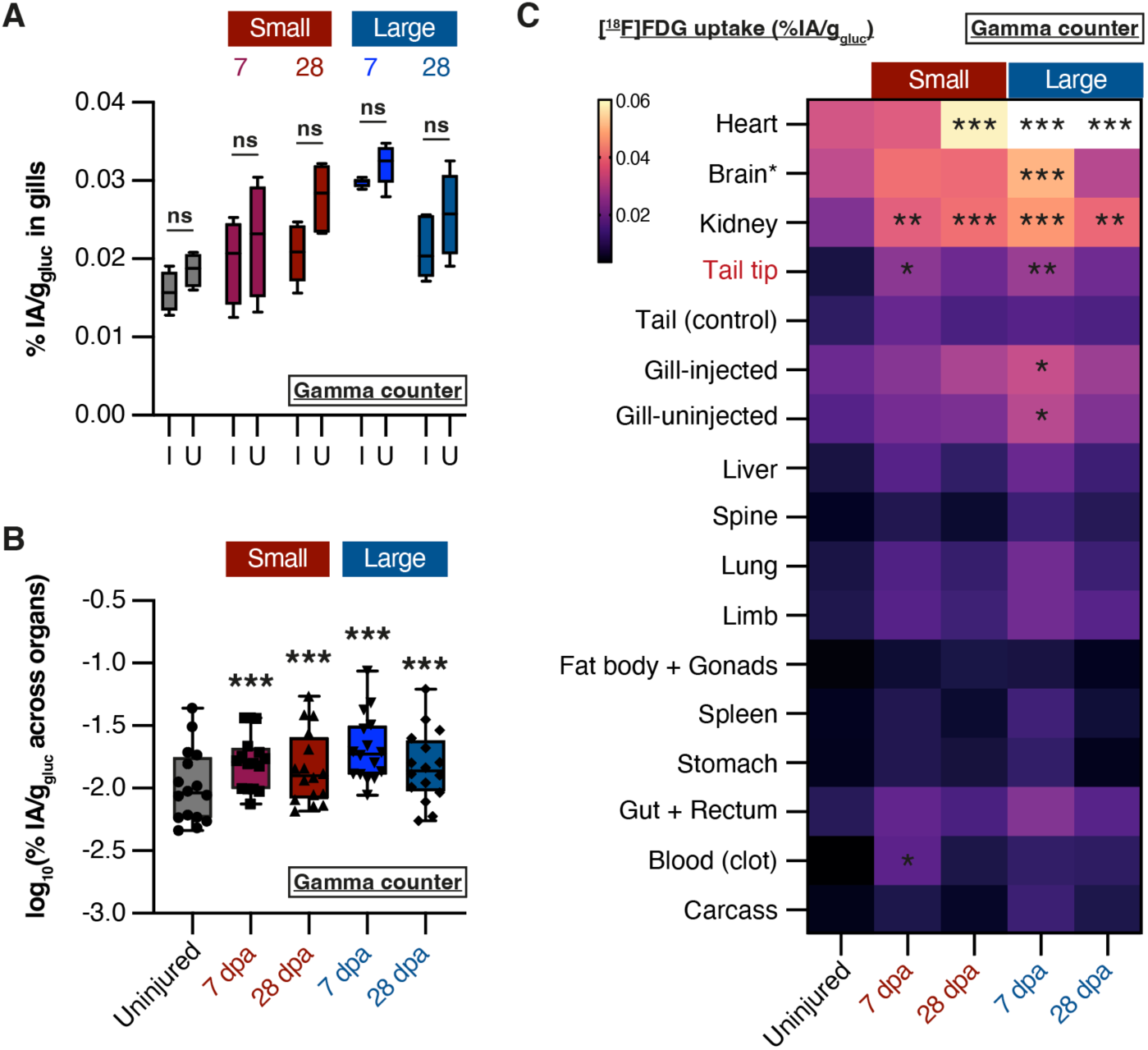
Gamma counter confirmations of PET quantification results. (A) Paired analysis of [^18^F]FDG uptake in injected gills (I) and uninjected (U) gills in each experimental group, measured by gamma counter. Data are displayed as the % of the total injected activity (IA), normalised to the weight of the tissue and multiplied by the blood glucose concentration of the animal. No significant difference (ns) was seen in any group. Multiple Wilcoxon tests, *p* > 0.05. *n* = 4 animals (control, small 7 dpa) or *n* = 5 animals (all other conditions). (B) Comparison of average %IA/g_gluc_ values across experimental groups (gamma counter). Each dot represents a different organ, and represents the mean value from all animals in that experimental group. The regenerating tail tip was not included in these data. Data were log-transformed to allow statistical comparison by 2-way ANOVA with Dunnett’s multiple comparisons. ***: *p* < 1×10^-12^. (C) Heatmap comparing %IA/g_gluc_ in individual organs in each experimental group (gamma counter). Displayed heatmap value is the mean %IA/g_gluc_ of all animals in that experimental group. Yellow colours indicate higher values, while purple colours indicate lower values. Statistical significance compared to uninjured controls was determined by 2-way ANOVA with Dunnett’s multiple comparisons. *: *p* < 0.05, *: *p* < 0.01, *: *p* < 1×10^-3^.

