## Supplementary Figures for "Injury size regulates glucose allocation locally and systemically during vertebrate tissue regeneration"

**Figure S1.**

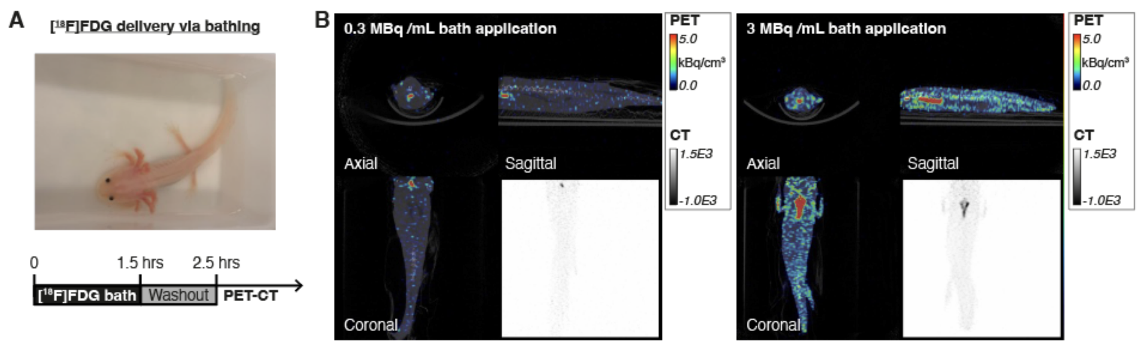


**Figure S1. Bathing is an inefficient method to deliver [$^{18}$F]FDG into axolotls in comparison to intravascular injection**

(A) Image of a regenerating axolotl in a water bath containing [$^{18}$F]FDG (above). Timeline summarising the experimental design (below). Only in this pilot experiment, PET imaging was paired with CT scanning to resolve skeletal structures (PET-CT). *n* = 3 animals.

(B) PET-CT images depicting tracer distribution after bathing axolotls for 90 mins in a [$^{18}$F]FDG-containing water bath with a radioactivity concentration of 0.3 MBq/mL (left) or 3 MBq/mL (right). The resulting total exposure was 50 MBq or 300 MBq respectively. Red indicates higher [$^{18}$F]FDG activity; blue indicates lower activity.

#### Figure S2.

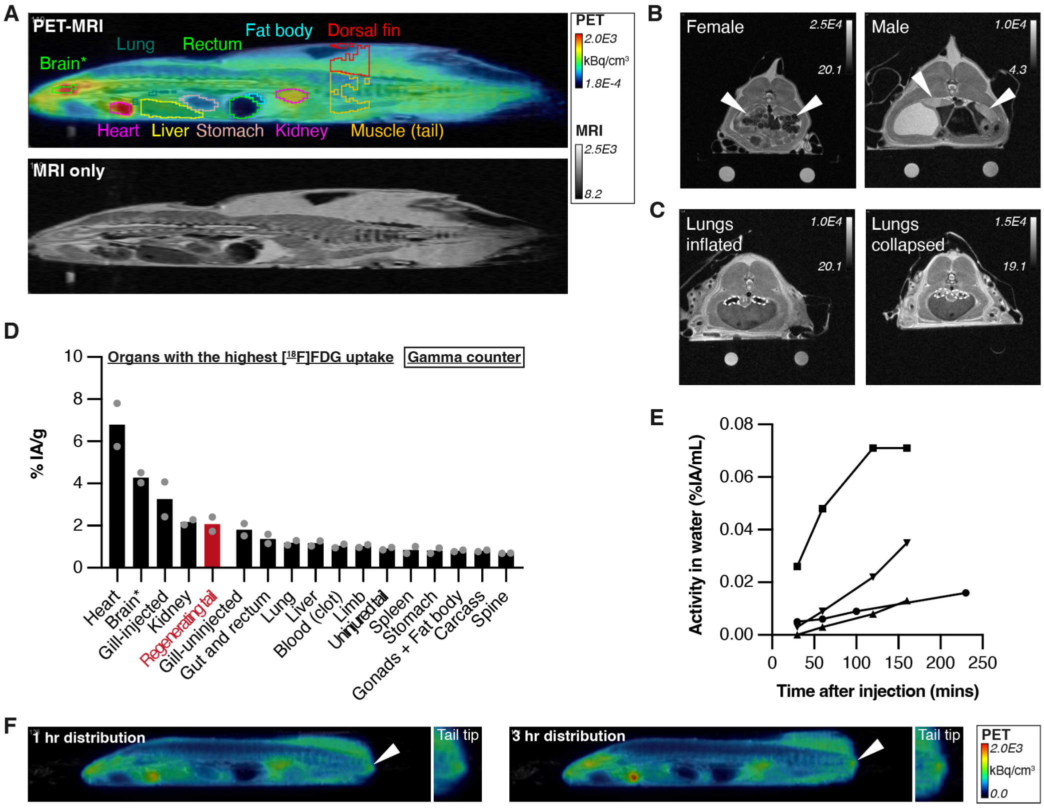


**Figure S2. Establishment of PET-MRI conditions for regenerating axolotls**

(A) A 14 cm axolotl 4 hours after injection with [$^{18}$F]FDG, depicted in sagittal view. The tail tip is not visible in this image plane. Top image shows PET signal overlaid on MRI data, with segmented organs indicated. Bottom image shows MRI data only.

(B) MRI images in axial view with reproductive structures arrowed.

(C) MRI images in axial view with lungs outlined.

(D) [$^{18}$F]FDG dose in selected organs 3 hours after injection, as determined by gamma counter measurements of dissected organs. Units are % injected activity (IA)/g. Arranged in decreasing order. The regenerating tail tip is highlighted in red. $n$ = 2 for pilot experiments. Brain*: forebrain and midbrain only.

(E) Estimation of [$^{18}$F]FDG leakage into the water bath (loss from the animal due to leakage or excretion). $n$ = 4 animals. IA: injected activity. The volume of the water bath was 400 mL.

(F) Representative PET images of the same axolotl imaged 1 and 3 hours after [$^{18}$F]FDG injection. Arrowheads indicates signal in the regenerating tail tip. Sagittal view, head pointing left.

### Figure S3.

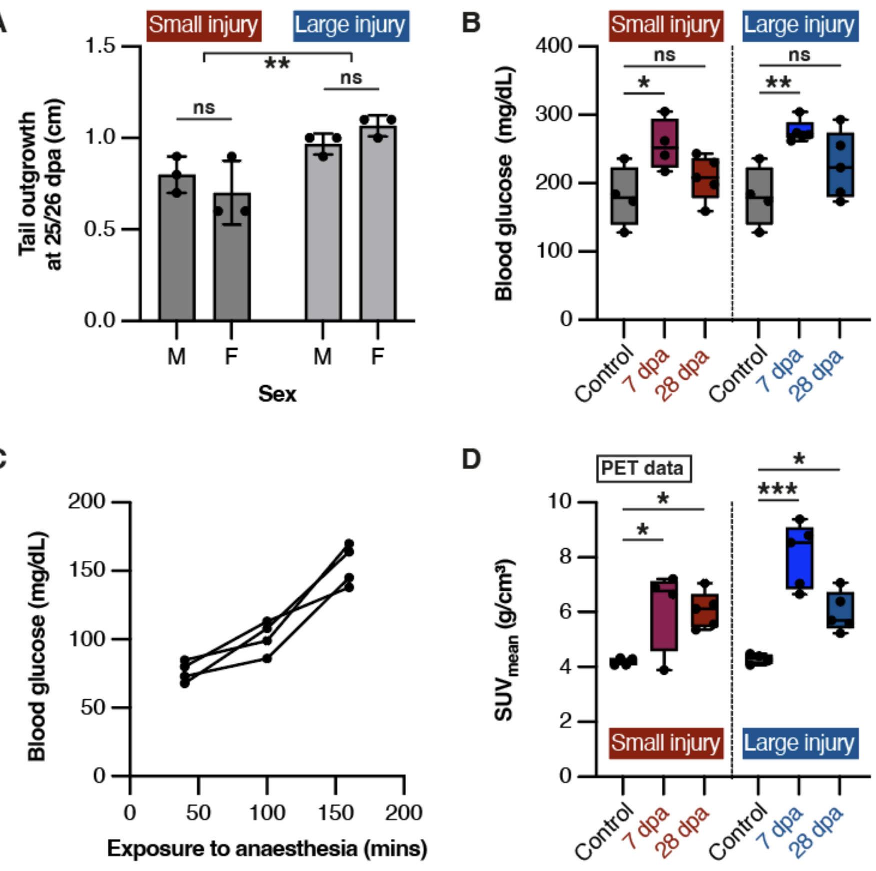


**Figure S3. Characterisation of tail outgrowth and blood glucose effects**

(A) Tail outgrowth at 25-26 days after small injury or large injury, separated by sex. Tails that received a large injury regenerated significantly more tissue than those that received a small injury (**: $p = 2.70\text{x}10^{-3}$). No significant different (ns) was observed between male and female animals ($p > 0.05$). Two-way ANOVA, $n = 3$ animals per group (12 animals total).

(B) Blood glucose levels in axolotls in each experimental cohort. One-way ANOVA followed by Šidák's multiple comparisons test. *: $p = 4.29\text{x}10^{-2}$, **: $p = 5.06\text{x}10^{-3}$.

(C) Blood glucose levels after increasing exposure to anaesthesia. Each line connects the datapoints from one animal. $n = 4$ animals.

(D) Quantification of [$^{18}$F]FDG uptake at the regenerating tail tip after small or large injury compared to the equivalent region of uninjured tails in control animals. SUV was not corrected for blood glucose concentration. One-way ANOVA, followed by a Holm-Šidák's multiple comparisons test. From left to right: $p = 2.08\text{x}10^{-2}$, $p = 1.97\text{x}10^{-2}$, $p = 1.10\text{x}10^{-5}$, $p = 3.82\text{x}10^{-2}$.

### Figure S4.

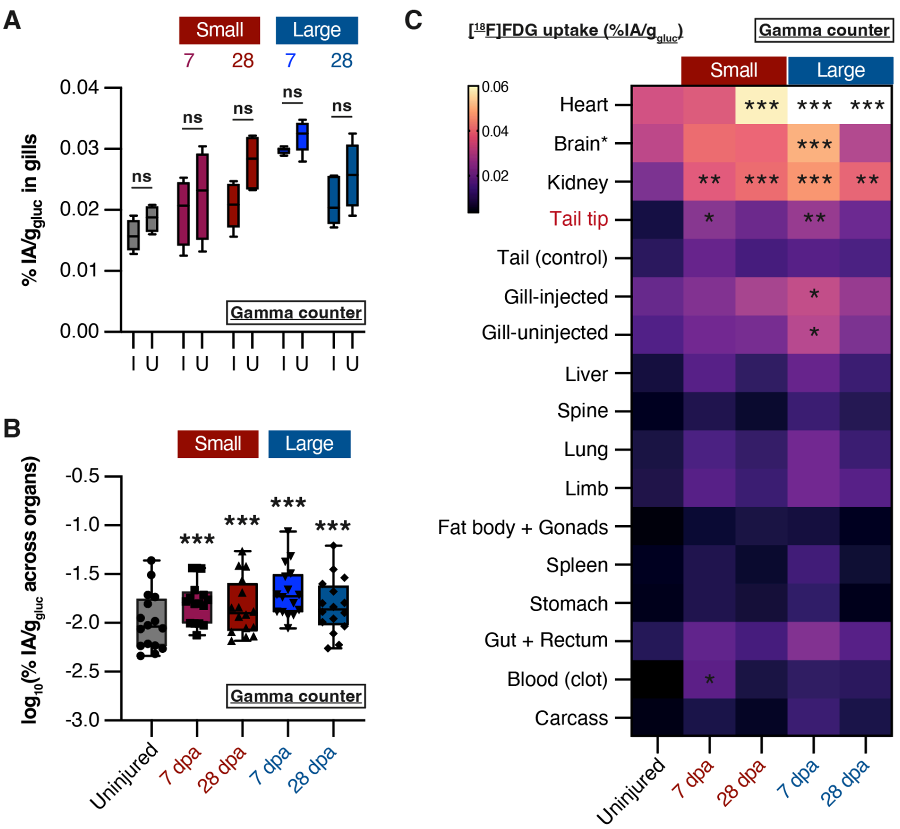


**Figure S4. Gamma counter confirmations of PET quantification results**

(A) Paired analysis of [$^{18}$F]FDG uptake in injected gills (I) and uninjected (U) gills in each experimental group, measured by gamma counter. Data are displayed as the % of the total injected activity (IA), normalised to the weight of the tissue and multiplied by the blood glucose concentration of the animal. No significant difference (ns) was seen in any group. Multiple Wilcoxon tests, $p > 0.05$. $n = 4$ animals (control, small 7 dpa) or $n = 5$ animals (all other conditions).

(B) Comparison of average %IA/$g_{gluc}$ values across experimental groups (gamma counter). Each dot represents a different organ, and represents the mean value from all animals in that experimental group. The regenerating tail tip was not included in these data. Data were log-transformed to allow statistical comparison by 2-way ANOVA with Dunnett's multiple comparisons. ***: $p < 1x10^{-12}$.

(C) Heatmap comparing %IA/$g_{gluc}$ in individual organs in each experimental group (gamma counter). Displayed heatmap value is the mean %IA/$g_{gluc}$ of all animals in that experimental group. Yellow colours indicate higher values, while purple colours indicate lower values. Statistical significance compared to uninjured controls was determined by 2-way ANOVA with Dunnett's multiple comparisons. *: $p < 0.05$, *: $p < 0.01$, *: $p < 1x10^{-3}$.
